# Two modes in the velocity statistics in cautious walks of laboratory rodents

**DOI:** 10.1101/2024.04.23.590757

**Authors:** I. S. Midzyanovskaya, A. A. Rebik, O. S. Idzhilova, V. V. Strelkov, N. L. Komarova, O. A. Chichigina

## Abstract

We have analyzed a large number of rodent tracks in open-field tests, in order to elucidate the statistics of their velocities. We found that the probability distribution of the absolute velocity of rodents can be approximated by a superposition of two Rayleigh distributions, with distinct characteristic velocities *v*_1_ and *v*_2_ with *v*_1_ *< v*_2_; this is in contrast to the single Rayleigh distribution for the velocity of a Brownian particle executing 2D random motion. We propose that the part of the distribution near the larger velocity, *v*_2_, characterizes rodents’ progressions in space, while the part near *v*_1_ describes other types of motion, such as lingering and body micromovements. We observed that the animals switched randomly between these two modes. While both velocities, *v*_1_ and *v*_2_, increase with age, their ratio, *v*_2_*/v*_1_, also grows with age, implying an increased efficacy of switches between the two modes in older animals. Since the existence of the modes is observed both in preweaned, blind pups and in older animals, it cannot be ascribed to foraging, but instead reflects risk assessment and proactive inhibition. We called such motion “cautious walks”. Statistical analysis of the data further revealed a biphasic decline in the velocity auto-correlation function, with two characteristic times, *τ*_*s*_ *< τ*_*l*_, where *τ*_*s*_ characterizes the width of velocity peaks, and *τ*_*l*_ is associated with the timing of the switches between progression and lingering. To describe the motion, we propose a stochastic model, which assumes the existence of two interfering processes: impulses to move that arrive at random times, and continuous deceleration. Its 2D Langevin-like equation has a damping coefficient that switches between two values, representing mode switching in rodents. Techniques developed here may be applicable for locomotion studies in a wide variety of contexts, as long as tracking data of sufficient resolution are available.

## I. Introduction

Laboratory rodents are among the most frequently used subjects of experimental neuroscience, and in particular, the studies of motor behavior [1, 2]. Quantifying rodent motor patterns is important for characterizing motor behaviors across diverse experimental scenarios, including human disease models, studies involving gene or circuit disruptions, investigations of post-induced brain or spinal cord injuries, and assessments of drug responses [3–7]. Motor assays, such as motion tracking, offer a costeffective and labor-efficient method to quantify different aspects of locomotor performance [8–10].

A number of statistical models of animal locomotion have been proposed. The motion of simpler species, such as invertebrates, has been successfully described by models based on the Brownian motion, see e.g. the reviews in [11, 12]. Evolutionary cephalization facilitated goaldirected locomotive strategies [13, 14]. A search for a randomly-distributed resources might include a Brownian strategy in mammalian and avian species [15–18], or its modifications such as correlated random walks [19– 21]. Models that contain Levy flights/ walks have been widely used to describe animal movement in the context of foraging, see e.g. [22–24] and reviews in [25, 26]. Many useful methods for studying velocity distributions and correlations have been developed in the field of microbiology [27–30].

In the mammalian locomotion models, animals are often assumed to move in order to find a rare reward. In a natural habitat, however, foraging is not necessarily the main driving force behind locomotion, as small animals are under continuous pressure to assess the risk of becoming prey [31]. Exercising caution is crucial for small animals to increase their chances of survival in environments where they face constant threats from predators. Evolution has shaped their behaviors and adaptations to minimize the risks associated with being preyed upon, which includes camouflage, sheltering, and nocturnal behavior.

Caution is also a natural feature of small animals’ spontaneous walks in a new environment. Animals evaluate potential environmental resources and dangers during stops, which usually punctuate locomotion [32] and are often used for reorientation. In case of danger, a stop may turn into a complete arrest (“freezing”) [9, 33], or it could be followed by a spontaneous flight to escape predation [34].

In this paper we used laboratory rodent (rat and mouse) experiments and mathematical modeling to create a statistical model of rodent walks, which incorporates deceleration to reflect caution, as a factor that modulates the statistics of the animals’ velocity. To dissociate reward-related foraging behavior from risk-avoiding environmental exploration, we analyzed short-term (of the order of minutes) tracking data from a large number of experimental observations of rodent walks in an open field. Our test subjects included adult rodents in a regular satiated state (i.e. not deprived of food/ water before the tests), as well as pre-weaned rodent pups of different ages, prior to the age of solid food ingestion. Therefore, in our experiments we assume that food search is not the leading motivation to move in a new environment. The tests were conducted by placing a single animal in an empty arena, in order to observe spontaneous (that is, not provoked by external events) locomotion in a stimulus-free space. We refer to this type of motion as “cautious walks”. In order to observe various stages of locomotion and spatial navigation, we included animals of different ages, to see if locomotor patterns exhibit any age-related features.

In this setting, we expect that the animal movement behavior is largely independent of the environment, but rather determined by the animal’s internal brain dynamics, at its different levels of motor control. Many authors have modeled the trajectories of this type of motion by using random walks, see e.g. [11, 32]. The question becomes, to what extent are these walks random and uncorrelated? The first step to answering this question is to analyse the velocity, rather that the trajectories of the animals. This is what we focus on in this paper.

We find that the statistics of individual instantaneous velocity of animals deviates from the classical Brownian particle in two ways: (1) The absolute velocity distributions for individual animals are not well described by a single Rayleigh distribution, but instead seem to contain two Rayleigh components with distinct characteristic velocities, and (2) the auto-correlation function of the velocity is characterized by two distinct correlations times, corresponding to short and long correlations, *τ*_*s*_ *< τ*_*l*_.

In order to describe these statistics, we created a model, where an individual animal accelerates according to a stationary pulse Poisson process. It is assumed to be subject to random pulses of acceleration, whose correlation time corresponds to the smaller value, *τ*_*s*_. On the other hand, deceleration is modelled as a consequence of “friction”, or “viscosity”, whose attenuation constant takes two values. The larger of the values corresponds to the mode of slow movements, which include the so called lingering, rearings, lateral head scans and short grooming, while the smaller one corresponds to the mode of “quasi-linear raids”, see [35]. The random switching between these two modes yields the second, large correlation time, *τ*_*l*_. The probability distribution of the absolute value of the velocity is given by the Rayleigh function for each of the two modes, while overall, the probability distribution has the form of a two-component Rayleigh function.

This model is an attempt to create a statistical description of rodent movements that is closely related to the experimental observations. Using this model, we propose causal relations between the different components of velocity control loops. Further, it can be used to study animal models of human motor disorders, emotional regulations of motor patterns, the degree of variation among animals in the same age-group, and also determine which parameters experience change as a result of development.

## II. Materials and methods

### A. Animals and track recording

The animals were housed in standard plastic cages, on a 12-h light/dark cycle, with food and water available *ad libitum*. All the rat pups included were of an outbred Wistar strain; adult rats were outbred Wistar (the in-house archive) or Long-Evans (the open source video archive, see below) males. The mice were of an outbred ICR strain. The numbers of animals in every age group are given in the Results section.

The rat dams were separated from their cage-mates since the last days of pregnancies. Mouse mothers were kept with the litter, the mate and, optionally, one more female cage-mate. The rodent litters were intact until the day of experiment. On the postnatal days 11 (mice, M11), 13 (rat, R13), 15 (rat, R15) or 17 (rat, R17) the dams were separated from their pups. The rodent nestlings were allowed to calm in their huddles for about 30 minutes. Then, the pups were gently grasped individually from the huddle periphery, and transported to the experimental chamber next door. Each pup was released in the center of an open field arena (black chamber 59*×*59 cm^2^, the same as reported in [35]), with a camera set 1m above. The pup’s placement was a start for video tracking which lasted for 120 seconds (*N* = 336), 300 seconds (*N* = 11), or 600 seconds (*N* = 14). The arena was cleaned after each individual session. All the pups were tested once, weighed after the experiments and returned to their mothers immediately. 14 rat litters and 4 mouse litters, of 6-14 pups, were enrolled. Body weights of the rodent pups are reported in table I.

**TABLE I.** The parameters of function (6) fitted for each animal group. The average body weight of the animals in each group is also provided.

| Parameter | M11 | R13 | R15 | R17 | R adults |
| --- | --- | --- | --- | --- | --- |
| $b$ | 0.76 | 0.78 | 0.70 | 0.79 | 0.89 |
| $\tau_s$, sec | 0.18 | 0.33 | 0.40 | 0.21 | 0.59 |
| $\tau_l$, sec | 12.23 | 9.95 | 5.50 | 3.40 | 14.27 |
| weights, g | $7.02 \pm 1.1$ | $22.7 \pm 4.0$ | $31.3 \pm 4.6$ | $36.4 \pm 7.2$ | 250-400 |

We used mouse and rat pups of a comparable developmental stage (M11 and R13; i.e., 2-3 days prior to eye opening), and two older cohorts (R15 and R17). The second rat pup cohort (R15) had started to open their eyelids, but had not yet acquired visual sensing [36], while the oldest pup group (R17) fully possessed the visual sensory inflow [36]. Therefore, the adult animals (R adult) and 17 day old pups (R17) were regular sighted rats, studied under normal day-light conditions, with fully available visual cues.

Video data of the adult rat group (R adult) consisted of 16 individual recordings made in a square (120*×*120 cm^2^) “open field” arena, according to the standard procedure: a rat was released at the arena center, with its back to the experimenter, and this was a start for video-tracking. The adult rats were males weighed 250-400g and aged 4-5 months. Additionally, an open video-archive of 20 Wistar rats performing an open field test (100*×*100 cm^2^ arena) was downloaded from example materials [37], and tracked as described below. The tracking data were pooled for analysis. The open field tests lasted 150 seconds (*N* = 20) or 300 seconds (*N* = 16) for adult rats (R adult).

Movement tracking was done offline. A tracker software (ToxTrack, the algorithm ToxID [38]) determined the body position by calculating the white mass center against the black background each 40ms. Briefly, animals were detected as bright moving objects using a threshold intensity value. The obtained objects or blobs were filtered by size to remove false-positive events.

The experiments were carried out in accordance with the National Institute’s of Health Guide for the Care and Use of Laboratory Animals; the experimental protocol was approved by the Institutional Animal Care Committee. All efforts were made to minimize animal discomfort.

## B. Trajectories

We digitized the motion of the rodents studied: M11, R13, R15, R17 and R adult. Typical trajectories are shown in figure 1, three for different types of animals, including a close-up (the bottom row).

**FIG. 1.**
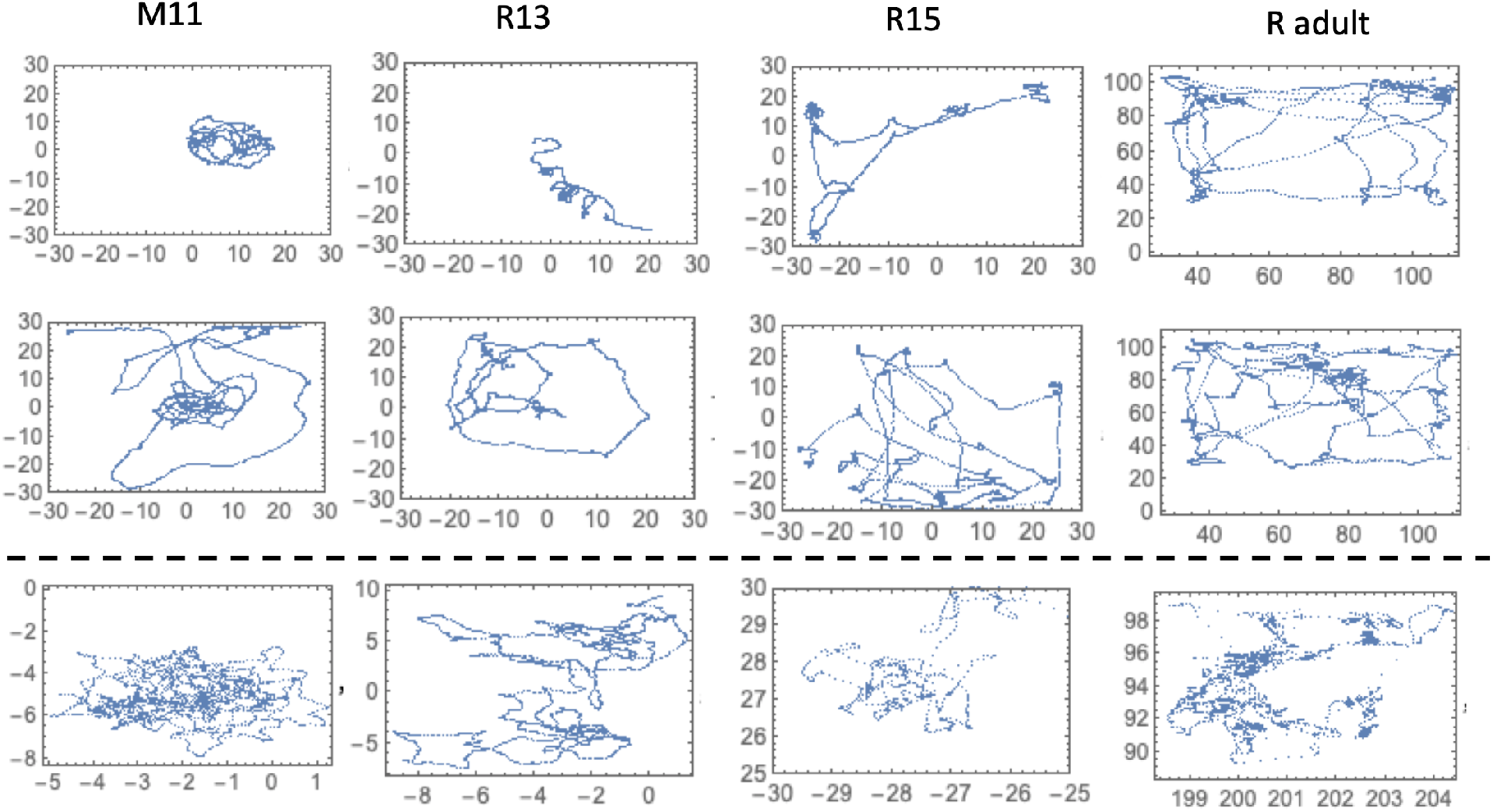
Trajectories corresponding to spontaneous walks of rodents: 11-day old mice and 13-day old, 15-day old rats, as well as adult rats). Three typical trajectories are presented for each animal group. The bottom row (below the dashed line) shows a close-up of a typical trajectory. The frames show distance in cm.

We note the presence of a large degree of variability in trajectory patterns, with some immature animals remaining within a few cm^2^ of their starting position, while others traversing the arena several times, and yet others spending time within a small area as well as engaging in longer runs. The blind walks of the youngest rat pups (R13) have been recently characterized as a superposition of localized walks and quasi-linear runs [35]. The trajectories obtained were typical for this type of tests, and included open center investigation, thigmotaxis (i.e. movements along walls), and localized walks during stoppings/lingering.

In all cases, visually, blind walk trajectories (M11,R13,R15) resemble those of Brownian motion, and support the suggestion that the motion is nearly isotropic.

## C. Velocity data collection

The data was obtained as coordinates of each animal at 0.04 second intervals. Denote the obtained 2D coordinates of the *i*th animal as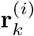, where *k* = 1, …, *N* are consecutive moments of time, *N* is the number of measurements in the time-series, and *i* = 1, …, *𝒩*, where *𝒩* is the number of pups in the sample. The absolute value of the instantaneous velocity at time *k*Δ*t* is numerically estimated as

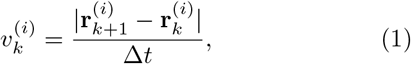

where Δ*t* is the duration of time between two measurements (0.04 sec). To gather the statistics of this quantity, all the data from the tracks were taken into account.

The auto-correlation function of the absolute velocity, Eq. (1), was calculated as follows. Define two sample means of the velocity:

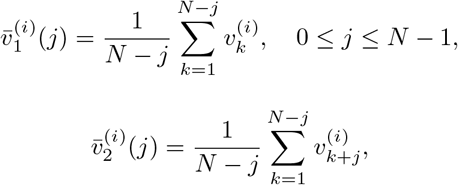

where again, the index *i* denotes the animal, index *k* the time-point, and *j* is time-shift. These quantities reflect the mean behavior of the different parts of the sample. Then, we define (see e.g. [39]) the auto-correlation function as

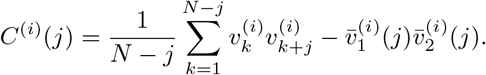

The normalized auto-correlation function (also called the auto-correlation coefficient) for pup *i* is then given by

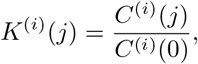

where the quantity *C*^(*i*)^(0) has the meaning of a sample variance. The average (over the rat pups) autocorrelation function for delay *τ* = *j*Δ*t* is calculated as

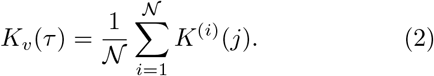

## III. Results

We investigated free walks of rodents in an open field. We digitized the motion of 5 groups of animals: 13-day old (*N* = 150) rat pups (R13), 15-day old (*N* = 159) rat pups (R15), 17-day old (*N* = 24) rat pups (R17), adult (*N* = 36) rats (R adult), and 11-day old (*N* = 33) mouse pups (M11). The young animals were of both sexes, since all pups in the litters were included. A part of the R13 dataset was analyzed in a recently published manuscript [35]. Adult rats were males, 4-5 months old, with body weights of approximately 250-400 g at the moment of recording. Typical trajectories are shown in figure 1, including a close-up (the bottom row). The digitized tracks have length between 120 and 600 sec for all the animals.

### A. The absolute value of the instantaneous velocity: the existence of two characteristic velocities

Figure 2 shows graphs of the absolute value of the instantaneous velocity of animals, where panel (a) shows motion over the interval of 100 seconds, while panel (b) zooms into a shorter interval of motion (5 sec). We observe the rugged appearance of these graphs with sharp accelerations and decelerations, suggesting the presence of short-time correlations. Despite a high degree of heterogeneity among the animals with respect to the speed of motion, some patterns can be discerned. For example, we can see in figure 2(b) that the average width of the high-velocity peaks is less than a second. Zooming out to panel (a), we further see that there are intervals of high peaks and intervals of low peaks of a typical duration of about a few seconds. Figure 3(a) illustrates some of these patterns further by plotting points of the trajectory in yellow if the instantaneous velocity was smaller than 1.5cm/sec, and in blue otherwise. We can see that the motion consists of an intermittent sequence of faster and slower regions. In figure 3(b) we present a typical graph of the absolute velocity as a function of time and denote the time intervals of slow motion as *ϑ*_11_, *ϑ*_12_, *ϑ*_13_, … and the intervals of fast motion as *ϑ*_21_, *ϑ*_22_, *ϑ*_23_,… One can see that there is a distinct difference between the regions of fast and slow motion. In the next sections we discuss this dichotomy in more detail.

**FIG. 2.**
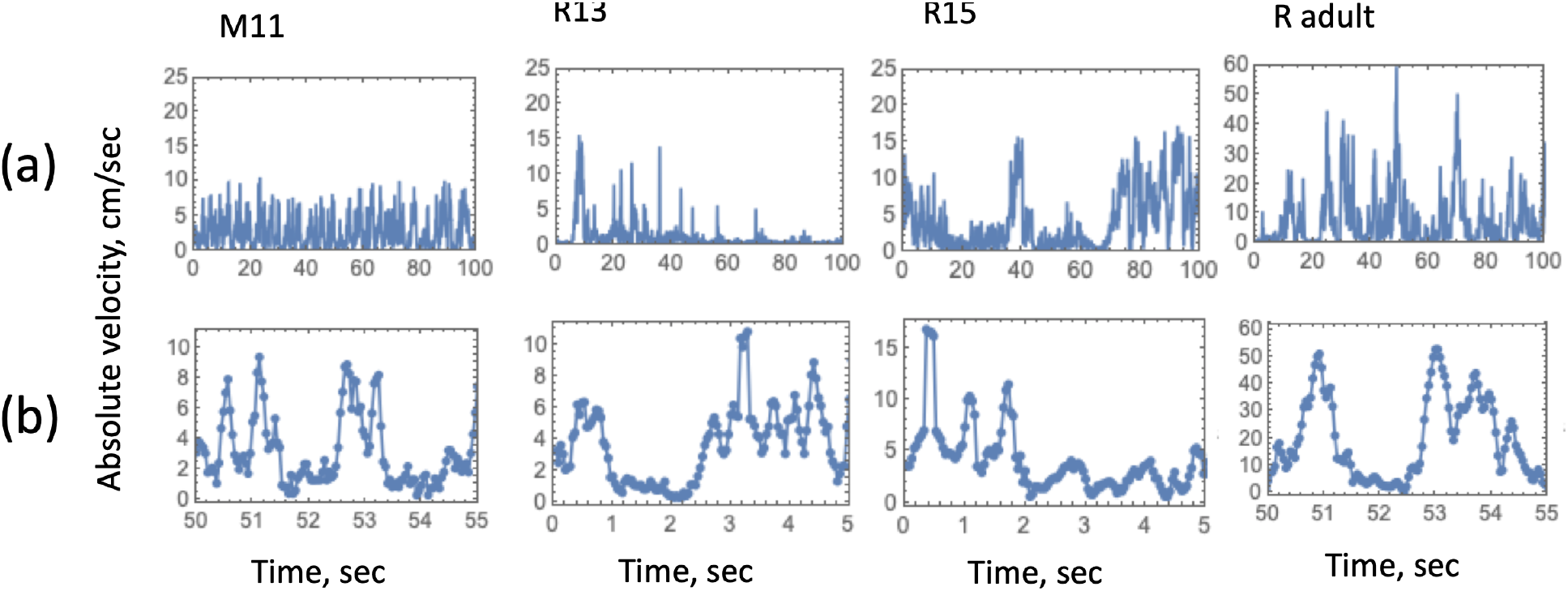
Typical graphs of the absolute value of the instantaneous velocity (Eq. (1)), plotted against time, for the four groups of animals. Please note the different range in the plots for adult rats. (a) Plotted over 100 seconds (note the different vertical scale for adult rats); (b) A close-up over the interval of 5 seconds.

**FIG. 3.**
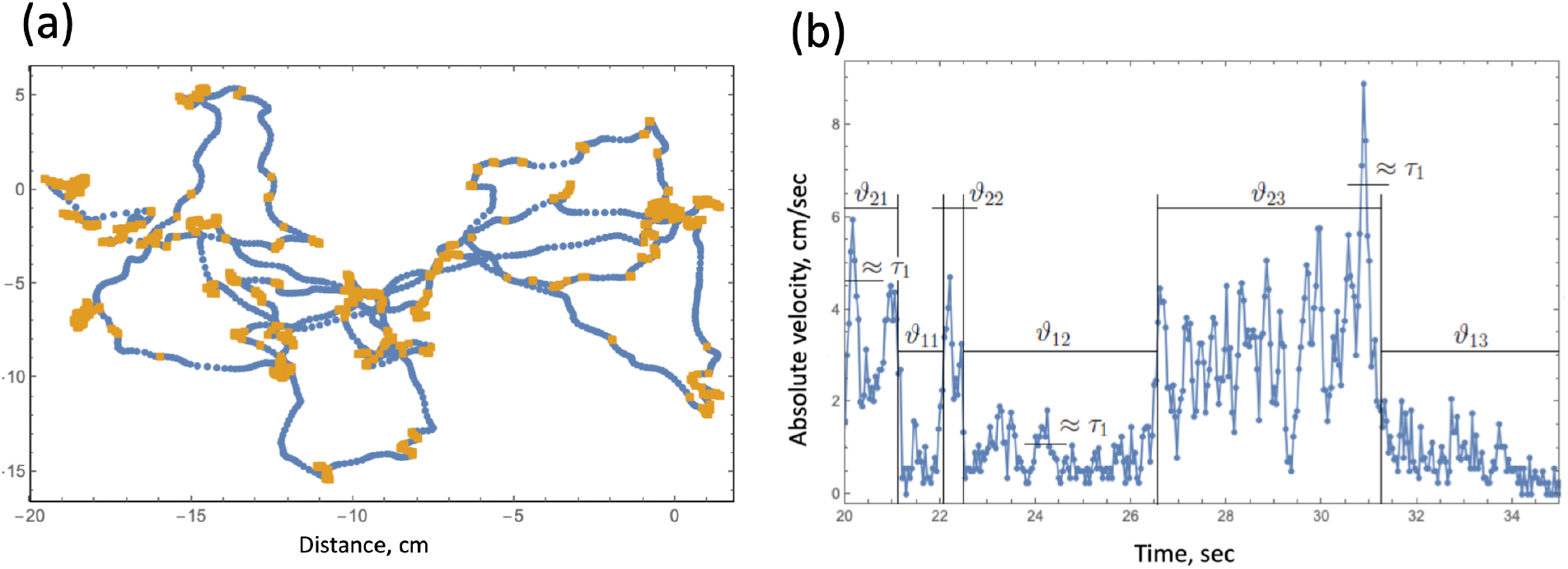
Velocity variation patterns. (a) An example of a typical trajectory of a 13-day old rat pup, where the yellow segments correspond to instantaneous velocities *<* 1.5 cm/sec, while blue segments to velocities ≥ 1.5cm/sec. (b) The two modes corresponding to strong and weak decelerations as dichotomous noise. *ϑ*_1*i*_ are the times of localized walks during lingerings and *ϑ*_2*i*_ the times of progressions.

Examples of the statistics of the absolute velocity are presented in figure 4, where we show histograms of the absolute velocity for individual animals. Such frequency vs velocity curves were fitted to several functions. First they were fitted to the function proportional to the Rayleigh distribution,

**FIG. 4.**
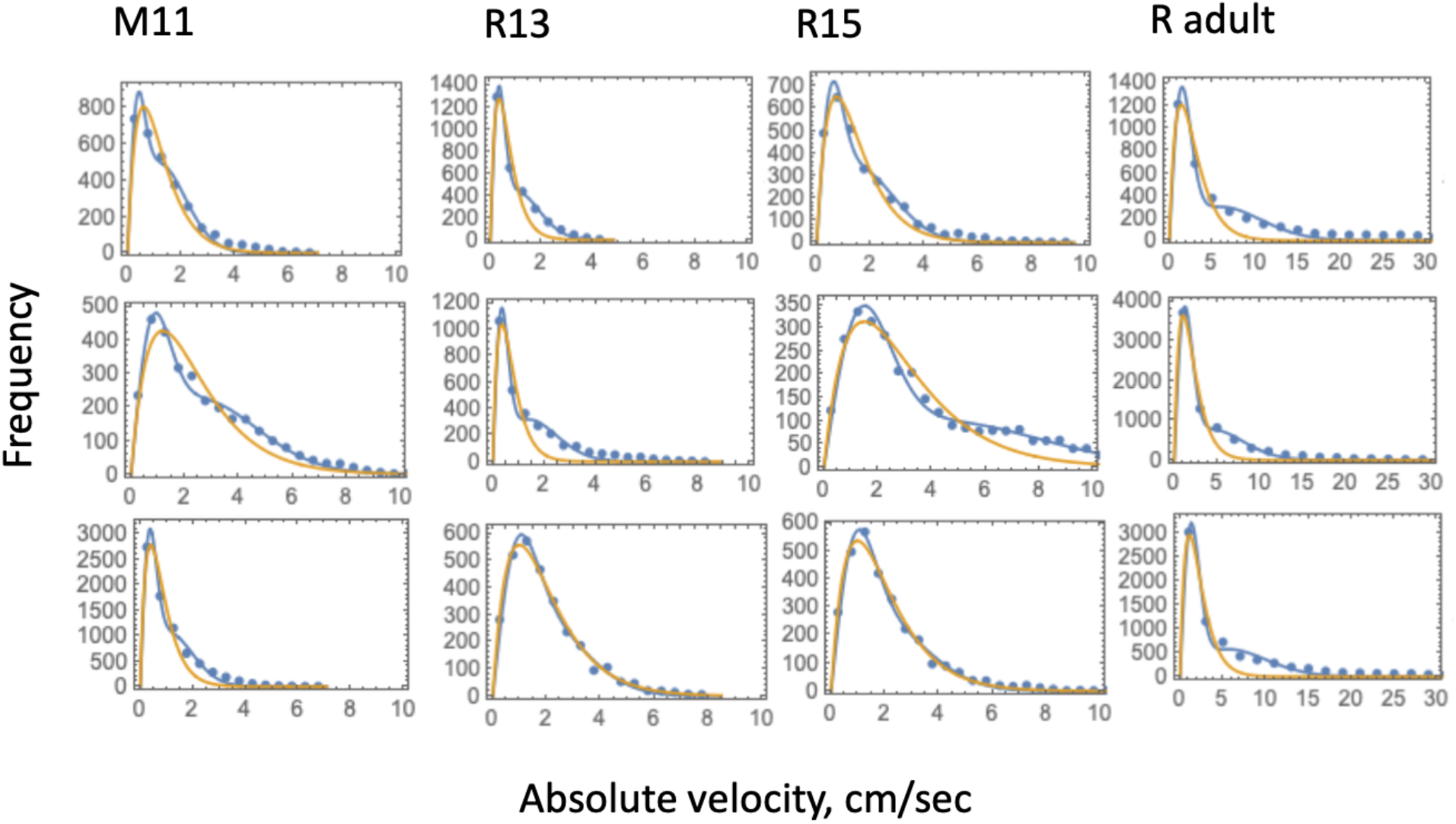
Sample graphs of the frequency of absolute values of the instantaneous velocity for the four groups of animals are presented by dots (with the bin size chosen to be 0.5cm/sec for all animal groups except for adult rats, where it was 2cm/sec). These functions are fitted by a functions (5) (yellow curves) and (4) (blue curves). The three graphs represent three examples of individual animals for each group.

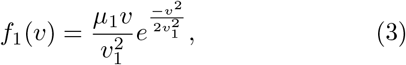

and then to a weighted sum of two Rayleigh distributions with characteristic velocities *v*_1_ *< v*_2_,

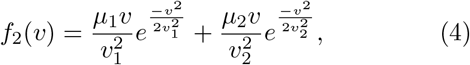

where *μ*_1_ and *μ*_1_ are the relative weights of slow and fast movements. It was apparent that in the great majority of the cases, the two-component Rayleigh distribution (function (4)) provided a better fit. The superiority of model (4) with respect to model (3) was further demonstrated by using the Akaike information criterion (AIC), see Appendix S1. The lower value of this criterion corresponds to the stronger modelling choice. Our calculations show that, despite having two additional parameters, function (4) is a stronger choice than function (3). Because of the two contributions with different characteristic velocities, the shape of this function better reflects a somewhat prolonged “shoulder” of the velocity distributions of individual animals.

In addition to the functions (3) and (4), we also tested a different two-parametric function,

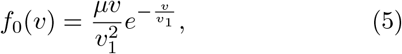

which is a Gamma-distribution with the shape parameter *k* = 2. The motivation behind this choice is to use a function that is zero at *v* = 0 and features a slower decay compared to the Rayleigh distribution. Again, this function’s AIC was compared to that for the two-component Rayleigh distribution. The results are shown in Figure 5 (see also Appendix S1). Again, the two-component distribution (function (4)) presented a better fit for the majority of cases, but here some interesting age-dependent patterns were observed, which we discuss next.

**FIG. 5.**
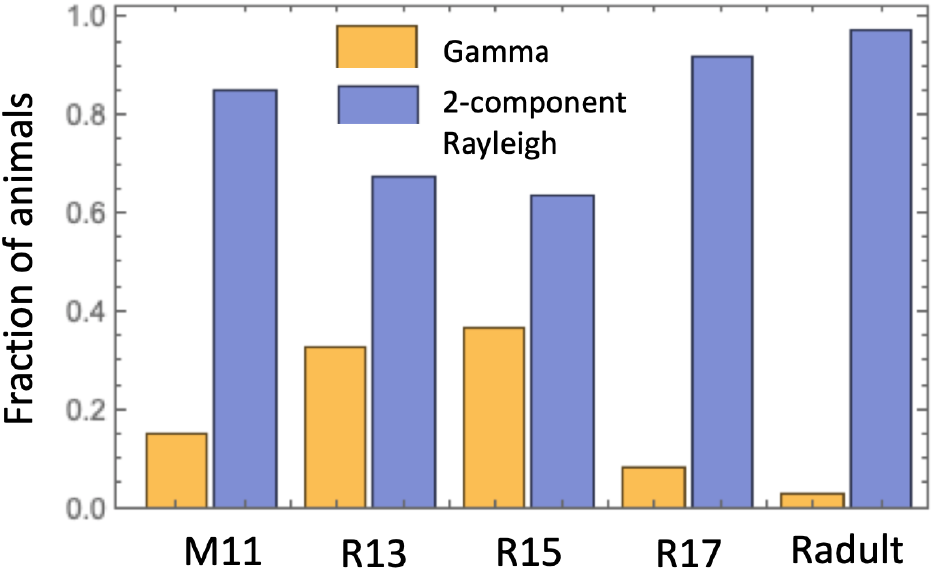
Model selection for the velocity distributions: comparing the Gamma-distribution (5) and a two-component Rayleigh distribution (4). Shown are the proportions of animals better modeled with the two-component Rayleigh distribution (blue) and better modeled with the Gamma-distribution (yellow).

For the youngest groups of 13-day old and 15-day old rat pups, the fit with function (5) resulted in a lower AIC for 32.7% and 36.5% respectively, suggesting the existence of a single characteristic velocity scale for those animals. The remaining 67.3% and 63.5% are better described by a two-component Rayleigh distribution. The mice pups on a comparable developmental stage (M11), were better described by the Gamma distribution in 15.1% cases. Among the 17-day old rat pups, 8.3% were better described by function (5), and 91.7% were characterized by two velocity parameters. Among the adult rats, 97.2% were better described by a two-component Rayleigh distribution.

Based on these data, we see a trend where as the animals get older, their velocity distributions tend to be described better by a two-scale Rayleigh distribution, with a sharp improvement reached by eye-sighted animals (R17 and R adult, vs M11, R13 and R15.) Presumably, the mature visual input helps in reorientation trials and thus in a better splitting of the modes.

Examples of fitting are presented in figure 4, where three instances of individual animals’ velocity distribution are plotted together with the best fitting function Gamma-distribution (5) (yellow curves) and a weighted sum of two Rayleigh distributions (4) (blue curves). The two terms in function (4) can be interpreted as two components of motion, progressions and lingerings.

Using the two-component Rayleigh function (4), we excluded 5% of cases with the largest mean squared error, and considered the dependence of *v*_2_ on *v*_1_ for the remaining values, see figure 6. The dependence is well described by a linear function with the proportionality coefficient between about 3 and 4, for all the animal groups.

**FIG. 6.**
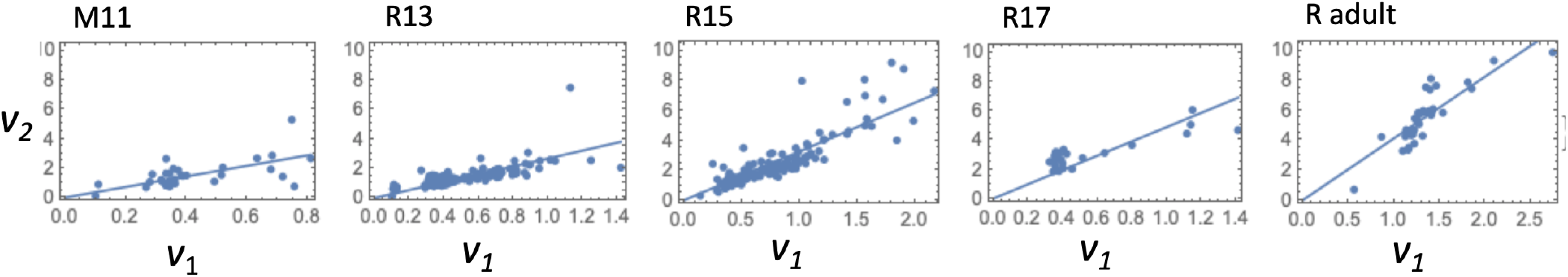
Best fitting characteristic velocity values, *v*_1_ and *v*_2_. Each dot corresponds to an individual animal; 5% of cases with the largest mean squared error are removed. The best fits with the linear function, *v*_2_ = *av*_1_ (a single-parametric fitting) are shown as straight lines. The values of the proportionality coefficient *a* are given by *a* = 3.58 for M11, and *a* = 2.67, 3.26, 4.87, and 4.13 for R13,R15, R17, and R adult respectively.

### B. Patterns of age-progression in the context of the two characteristic velocities

Figure 7 shows the means of the two characteristic velocity values for each group of animals. There is a clear age progression for the values of *v*_1_ and *v*_2_, which significantly differ from one another (see Appendix S1).

**FIG. 7.**
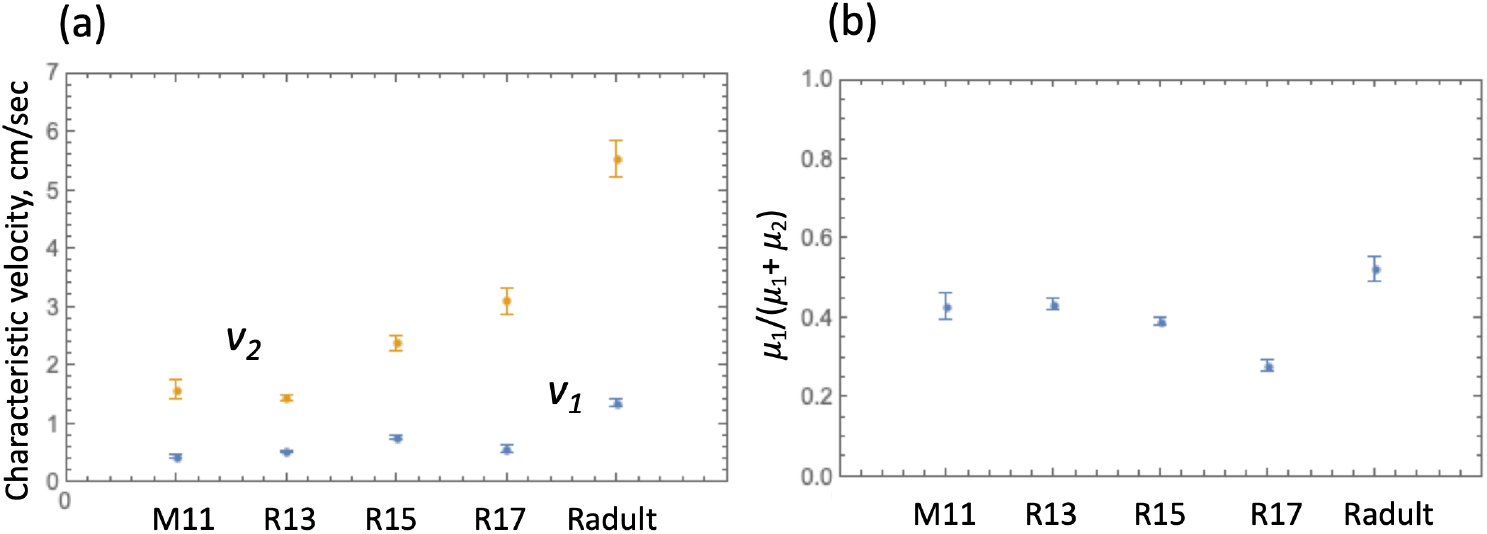
Characteristics of motion for different groups of animals. (a) The means, and standard errors of the means, for the best fitting values for *v*_1_ (blue symbols) and *v*_2_ (yellow symbols), for each group of animals. All the means for *v*_2_ are different among the groups (*p <* 0.05 by T-test). (b) The values of *μ*_1_*/*(*μ*_1_ + *μ*_2_), that is, the relative contribution of small motions, for all animal groups.

Interestingly, the relative weight of slow movements, *μ*_1_*/*(*μ*_1_ +*μ*_2_), seems to be stable during ontogeny and between the two rodent species: the micromovements constitute as much as a half of registered movements (Figure 7(b)). The functional blindness of the previsual pups (M11, R13 and R15) also does not affect the locomotor ratio studied. During the free walks inside the empty test arenas, possible navigation efforts of the animals appear to influence trajectories, but not the studied velocity dynamics and statistics.

### C. Two modes in the velocity auto-correlation function

We have calculated the normalized velocity autocorrelation function, or auto-correlation coefficient (see Section II C). Figure 8 presents the results, where the blue lines (experimental data) show a distinct bi-phasic decline. These curves were fitted with the function

**FIG. 8.**
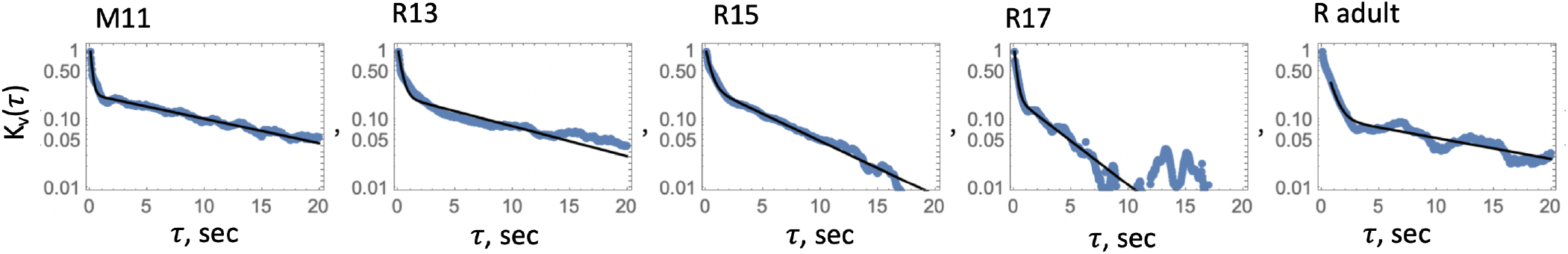
The velocity correlation function calculated by the method of Section II C (blue lines) together with the best fitting function (6), for the five groups of animals.

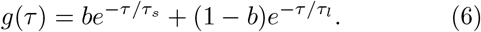

The black lines in figure 8 represent the best fit results for function (6). The values of *b, τ*_*s*_ and *τ*_*l*_ thus obtained are summarized in Table I.

The existence of two modes in the velocity autocorrelation function inspired a model that is described next. In the model, the two correlation times are understood as corresponding to two mechanisms of the speed regulation: acceleration and deceleration.

### D. Stochastic modeling of rodent walks

We have formulated a stochastic model that describes the motion of rodents, which is based on the observations reported above. If the rodent motion was completely random,[40] then its velocity projections would be distributed normally, and the absolute velocity would be described by the Rayleigh distribution [41]. This is a direct consequence of the Langevin equation with a constant noise intensity and a constant attenuation coefficient. In this case, the velocity auto-correlation function would be exponential with the characteristic time inversely proportional to the attenuation coefficient. This however is not what we observed.

Indeed, our data analysis demonstrated that the velocity is distributed according to a two-component Rayleigh function with two characteristic velocities; a deviation from the Rayleigh distribution indicates the presence of a correlation in acceleration [42, 43]. Further, the velocity auto-correlation functions for the rodents in our experiments are characterized by not just one, but two decay times.

In order to proceed with the mathematical model, we further note that the time-dependence of the instantaneous velocity presents as a sequence of peaks, whose width is similar to the smaller of the two correlation times (*τ*_*s*_ in Table I), see e.g. figure 3(b). Moreover, we observe the existence of two types of regions in the time-series, those with relatively low peaks and those with relatively high peaks. The duration of those regions varies, but is similar, roughly, to the larger of the two correlation times (*τ*_*l*_ in Table I). The envelope of the velocity vs time graph resembles a dichotomous process, with a sequential alternation of higher and lower horizontal regions (figure 3(b)). Based on the idea of the previously reported “gears” that were observed in the adult rodent locomotion [44], we propose the existence of two distinct modes in our experimental system, which correspond to (on average) faster and slower movements. The slower mode corresponds to behaviors such as lingering, lateral head scans, rearings, and short-term grooming on the one hand, and the faster mode to quasi-linear raids. An animal “turns on” one of the modes and moves accordingly, until it switches to the other mode, and so on. Those random switches are hypothesized to give rise to the larger correlation time, *τ*_*l*_.

To construct a stochastic differential equation for the velocity and to obtain probability distribution function (4), it is convenient to split the change of the velocity into acceleration and deceleration,

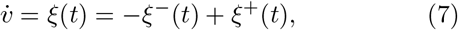

and construct a model for each of these two processes.

### a. Deceleration

In the absence of environmental cues, the switching between the slower and the faster modes is modeled as transitions in a two-level system [45], or dichotomous noise [41].

Let *ϑ*_1*i*_ be the times of slower movements during lingerings and *ϑ*_2*i*_ the times of faster walks, see Fig. 3(b). These two modes correspond to strong 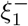 and weak 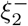 decelerations, respectively. For example, in Figure 3(b), we observe faster motion during the first second, that is, *ϑ*_21_ ≈ 1 sec, and then slower motion for about one second (*ϑ*_11_ ≈ 1 sec), then *ϑ*_22_ ≈ 0.5 sec, *ϑ*_12_ ≈ 4 sec, and so on. Transitions from one mode to another are random and uncorrelated. As a result, *ϑ*_1*i*_ and *ϑ*_2*i*_ are independent exponentially distributed random variables similar to the assumption in [17]; we denote their averages as *(ϑ*_1*i*_) = *T*_1_ and *(ϑ*_2*i*_) = *T*_2_. The ratio of these times is determined by the ratio of relative weights of slow and fast movements in experimental distribution (4):

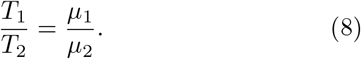

The correlation function of the deceleration is a correlation function of the Markovian process in a two-level (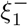 and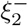) system [45],

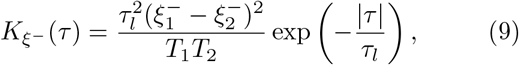

where the correlation time of deceleration (*τ*_*l*_ in Table I) is

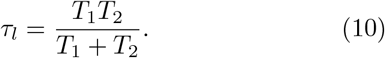

### b. Acceleration

Next we turn to the description of acceleration. We propose that locomotion of animals in a homogeneous environment corresponds to Poisson statistics of acceleration impulses:

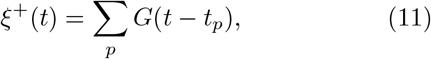

where *t*_*p*_ are random start times of the pulses and

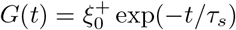

is the shape of the pulses, where 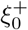 (cm/sec^2^) is an amplitude of the acceleration and *τ*_*s*_ is a characteristic width of each pulse.

As demonstrated below, this shape corresponds to an exponential auto-correlation function of acceleration with the characteristic time given by *τ*_*s*_ from Table I. The pulses do not correlate with one another, at least in the absence of environmental cues. The correlation of acceleration occurs due to the correlation of the pulse with itself: the time-lapses between two successive pulses, or waiting times (WT) Δ_*p*_ = *t*_*p*_ − *t*_*p*−1_, are distributed exponentially with the mean (Δ) = *T* . The width of the pulses *τ*_*s*_ is assumed to be small compared to the long correlation time: *τ*_*s*_ *« τ*_*l*_. The average of the pulse process is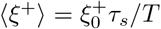.

The pulse process *ξ*^+^(*t*) can be represented using a sequence of *δ*-shape pulses or a point process [46, 47]

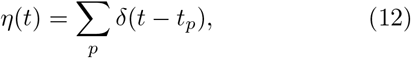

such that

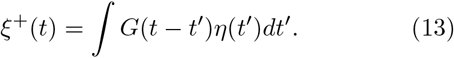

The spectral density of the process 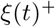 can be expressed as follows [48–50]:

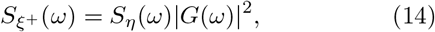

where *G*(*ω*) is the Fourier transform of the shape function *G*(*t*) and

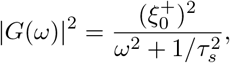

and the spectral density of the sequence of *δ*-shape pulses is

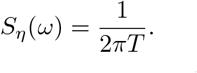

Substituting these expressions into (14) and making the Fourier transform in accordance with the Wiener-Khinchin theorem [51, 52] we obtain the correlation function of the acceleration

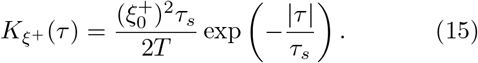

Exponential acceleration correlation is often used to describe animal movement [53].

### c. Correlation function of the velocity

We assume that the acceleration and deceleration are independent of each other, therefore the correlation function of the acceleration-deceleration is a weighted sum of the two functions, (9) and (15),

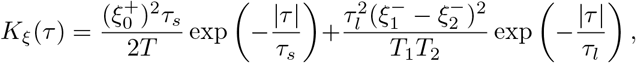

which is a sum of two exponents. The correlation function of the velocity can be estimated using the following equation [49, 50]:

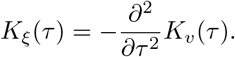

These transformations, related to the calculation of derivatives or integration, do not change the structure of the function consisting of exponents. As a result, the correlation function of the velocity also contains two exponents:

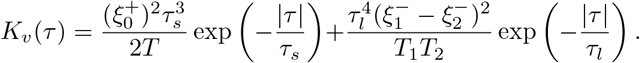

Comparing the ratio of the two prefactors in this theoretical correlation function with the ratio in the experimental one given by Eq. (6), we find:

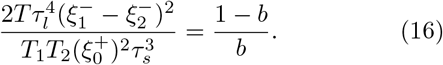

### d. Langevin equation

The conventional method of studying dynamics of a fluctuating parameter is to write a stochastic differential equation in accordance with the law of motion, and to obtain the probability distribution as a result. Our situation is reversed. We have a PDF deduced from experimental observations (4), and want to reconstruct the underlying law of motion. Each of the two Rayleigh distributions can be obtained from the standard Langevin equation for a two-dimensional Brownian motion with a damping coefficient *γ* and the noise intensity

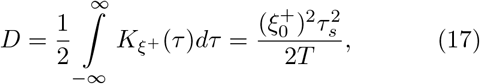

where the correlation function of the short-correlated noise *ξ*^+^ is given by Eq. (15) . We have two Rayleigh distributions corresponding to the two modes. We suggest that, for each of the two modes, the noise is the same, but the damping coefficient is different and given by *γ*_*i*_, where *i* = 1, 2 is an indicator of the mode. This coefficient changes as a dichotomous process. Substituting acceleration and deceleration into (7), we obtain the following Langevin equation corresponding to the PDF given in Eq. (4):

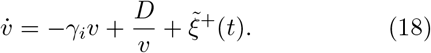

Here the second term comes from a two-dimensional Wiener process, or a Bessel process. The noise 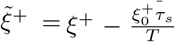 has a zero average and can be considered a white noise, since *τ*_*s*_ *« τ*_*l*_. We added a constant value 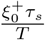 to the deceleration, to compensate the average acceleration. The deceleration also has a part proportional to the velocity, −*γ*_*i*_*v*. The stationary PDF corresponding to this equation is a weighted sum of two Rayleigh distributions. Comparing it with Eq. (4) we have:

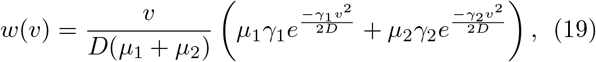

where 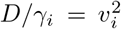 and *i* = 1, 2 is an indicator of the mode.

### e. The values of theoretical parameters

Using Eqs. (8) and (10), we obtain the average duration of each of the modes,

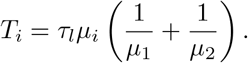

The noise intensity can be obtained from the ratio of the prefactors (16) in the velocity correlation function,

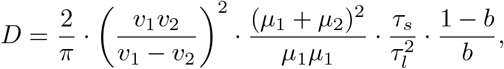

where we used (17). The changing parts of the deceleration are taken as 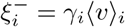, where the average velocity for each mode is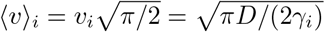, and specific parameter values can be found in Table I.

### f. A mechanistic analogy

In each of the two modes described here, a part of the graph looks similar to the graph of the velocity of a Brownian particle for two-dimensional motion and gives us the opportunity to describe the motion using Langevin-like Eq. (18). As a result, the rodent can be modeled as a Brownian particle with the mass and size, which are constant but different for each mode (see Appendix S1).

## IV. Discussion

Locomotor tests in laboratory rodents are part of experimental routine worldwide. One of the basic ways to observe animals’ behavior is to track their movements [54]. Since motion tracking software became available, animal locomotion has been digitized routinely. Mainly, it is done by calculating the instant body centroid coordinates against the background. This approach provides trajectories with a relatively high resolution.

In this paper, we have digitized and analyzed a large number of rodent tracks in open-field tests, in order to elucidate the dynamics of the slowest locomotor grade, which is pronounced in a novel environment. We called this type of motion “cautious walks”. We found that the probability distribution of the absolute instantaneous velocity of rodents can be approximated by a superposition of two Rayleigh distributions, which are characterized by distinct characteristic velocities *v*_1_ and *v*_2_ with *v*_1_ *< v*_2_; this is in contrast to the velocity of a Brownian particle executing random motion, which obeys a single Rayleigh distribution. The two contributions come with approximately equal weights. We propose that the part of the distribution associated with the larger of the two velocities characterizes rodents’ progressions in space, while the part of the distribution near the smaller velocity describes a range of other types of motions, such as lingering with closing steps [55, 56], body micromovements: lateral head scans, torso switches, etc [57].

Information on both larger scale motions (locomotion) and smaller movements is likely contained in many laboratory studies of movements that use motion tracking [58, 59]. The most common frontal view used to monitor test animals will pick up small movements of the head and the torso. Therefore, by necessity, the obtained signal contains different types of movement, such as forward progressions, lingering episodes, and vertical activity. Oscillatory-like lateral head scanning movements appear during lingering, and represent common behavioral elements, associated with assessing the environmental information in rats [57]. Vertical activity (rearings) are often used as a quantitative measure for investigatory drive in laboratory rodents [3, 4]. Therefore, the slow episodes during displacements are linked with orientation and decision making, and the corresponding proactive (“made in advance”) deceleration is an adaptive feature. The assumed proactive deceleration, as an essential feature of “cautious walks”, contrasts with the assumption in common foraging models, that a raid ends if (and when) the animal obtains a cue from the environment (see e.g. [53]).

Since in our experiments the rodent pups (below the age of solid food ingestion) demonstrated velocity patterns similar to those in adult animals (figures 4, 6, 8), we could not ascribe the stops or lingering to putative foraging attempts. As the decelerated episodes were characterized by the lateral head scans [57] and used for reorientation [32], we hypothesize that they reflect the process of risk assessment, necessary for prey animals in the wild. One might attribute the animals’ deceleration (i.e, action-stopping) behavior to the so called proactive inhibition [60–62], a condition where inhibition is strongly facilitated.

Apart from the existence of two characteristic velocities, statistical analysis of the data also revealed a biphasic decline in the velocity auto-correlation function, with two characteristic times, *τ*_*s*_ *< τ*_*l*_. The characteristic time *τ*_*s*_ characterizes the width of velocity peaks, and *τ*_*l*_ is associated with the timing of the switches between progression and lingering. These patterns inspired a stochastic model that is compatible with the observations of the rodent cautious walks. A central observation is that the animals slow down in order to re-evaluate the environment, with its potential danger and rewards (these stops might further develop into fear- or startle-induced behavioral arrests [33], or present as just a punctuation in the ongoing locomotor activity [32]). To describe this, we assumed the existence of two interfering processes: in addition to the impulses to move arriving at random times, continuous deceleration must be taken into account. This leads to the pulse structure of the dependence of velocity on time (see figure 2).

Our model can be contrasted with the “race model”, where during task discrimination trials, the execution and inhibition of an action are often presented as two concurrent motivations (reviewed in [63, 64]). This race model approach implies a competition of “stop” and “go” processes, and the winner determines the final action. Instead, our theory suggests a summation of two opposite effects acting upon the locomotor output, which results in acceleration and deceleration impulses.

The main finding reported here is the existence of two different modes characterising cautious rodent walks. Two methodological components are necessary to observe this behavior. (1) The data corresponding to this type of motion are made available by the technology that is currently used, which provides a frontal view on the dorsal surface of an animal and involves a subsequent calculation of the centroids, for each frame. Previously, researchers used manual or semi-automated ways to assess locomotion, where the experimental arenas were divided into (virtual) zones and border crossings were counted over time. As a result, no detailed information on micro-movements was available. In the present setting, while we face a challenge of filtering or separating the components, we also gain a potentially new source of information. (2) It is essential that the statistical analysis of animal velocity is performed on an individual basis. Velocity distributions were constructed and analyzed separately for individual animals. Pooling animals together and analyzing the ensemble behavior produce different statistical results, which are subject of a future work.

Having identified two different models of motion with different characteristic velocities, we can analyze some of the implications of this finding. A surprising observation was that the two characteristic velocities, *v*_1_ and *v*_2_, while quite variable among individual animals, are roughly proportional to each other. Our study provides an estimate for the ratio between *v*_2_ and *v*_1_ of about 2 to 5. This means that the energy cost for regular locomotion is an order of magnitude higher than what is needed for the micro-movements. Presumably, this estimate will be shifted by neuromuscular/neurodegenerative disorders, drugs affecting the vigilance state, etc. Further studies are needed to pursue these ideas.

The proportionality of *v*_1_ and *v*_2_ might also point to the idea of slow local movements as a precursor of major locomotor acts [55, 56]. An analogy may be drawn between these patterns and other cases involving smaller and larger motions, such as movements of athletes prior to start of run trials (“warming-up”), or motor behavior of patients recovering from lesion-induced akinesia [56], as well as comatose or deeply anaesthetized patients; in such cases, minor motor acts appear before gross locomotor activity. The same is known for the ontogenetic flow of motor skills: local motions of limbs precede its ability for large-scale movements [55].

Both characteristic velocities, *v*_1_ and *v*_2_, have been found to increase with age. This can be naturally attributed to a growing muscular power, as the simplest explanation. It is interesting that the ratio between the two velocities, *a* = *v*_2_*/v*_1_, also grows (albeit relatively slowly) with age, which implies an increased efficacy of switches between *v*_1_ and *v*_2_ in older animals. In about 1/3 of the young blind rats (R13, R15), the velocity distributions appeared to be blended, with poorly discriminated peaks for *v*_1_ and *v*_2_ (Figure 5). For older animals, we observe a clearer separation between the two models of motion. The separation of locomotor modes in more mature animals might therefore serve as a developmental milestone, marking the activation of a stress-response system [65], which initiates escape reactions.

Techniques developed here may be applicable for locomotion studies in a wider range of contexts. For example, by analyzing data archives accumulated for various drug-affected and/or pathological conditions, one may gain new insights into the mechanisms of a potentiation of gross locomotor acts by micro-movements, in a variety of physiological states. Such an empirical tool to estimate the action of inhibitory control over body-micromovements could be particularly important for modeling developmental disorders [66–68], ADHD, [69–71], Parkinson disease [72–74], etc.

## V. Acknowledgments

The authors would like to thank Prof. Vladimir V Raevsky and Dr. Marina L Pigareva for their valuable scientific discussion of ontogeny of laboratory rodents.

## Appendix A Further details of the statistical analysis of the models

Figure 9 shows three examples from each animal group, where the velocity distributions were fitted with functions *f*_1_(*v*) given by Eq. (3) of the main text (a single-Rayleigh distribution, yellow) and *f*_2_(*v*) given by Eq. (4) of the main text (a two-component Rayleigh distribution, blue).

**FIG. 9.**
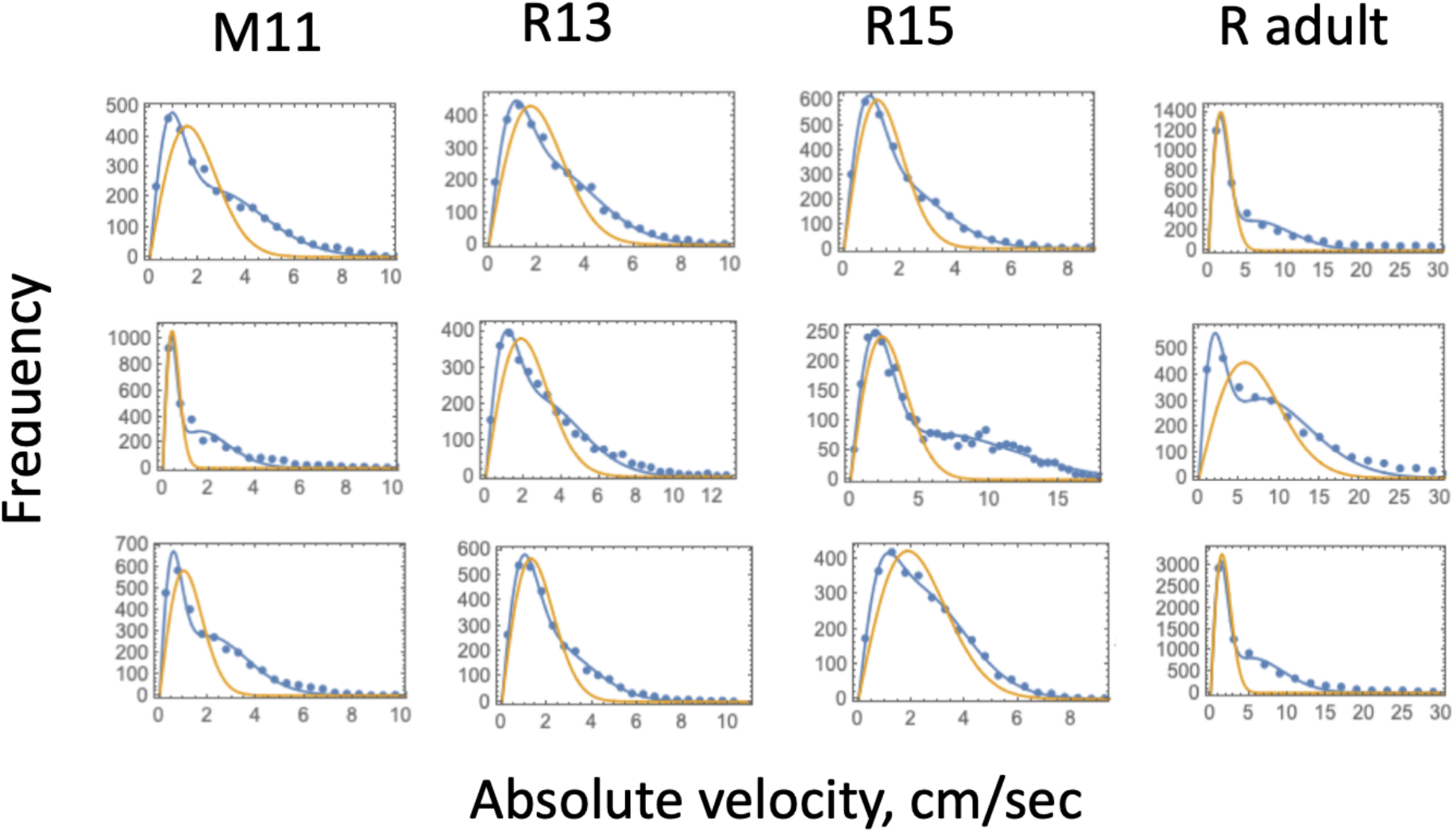
Graphs of the frequency of absolute values of the instantaneous velocity for four groups of animals are presented by dots (with the bin size chosen to be 0.5cm/sec for all animal groups except for adult rats, where it was 2cm/sec). These functions are fitted by a functions (3) of the main text (yellow curves) and (4) of the main text (blue curves). The three graphs represent three examples of individual animals for each group.

We used the Akaike Information Criterion (AIC) to check the relative power of models (3) and (4) of the main text in fitting the individual probability distributions of absolute instantaneous velocity. This statistic was calculated for all fits and denoted by *AIC*_1_ and *AIC*_2_, respectively. The better model corresponds to a lower value of the AIC. Figure 10 plots the difference, *AIC*_2_ −*AIC*_1_, for each of the velocity distributions; each dot represents a single animal. We can see that for all the animal groups, the overwhelming majority of velocity distribution fits yield *AIC*_2_ *< AIC*_1_.

**FIG. 10.**
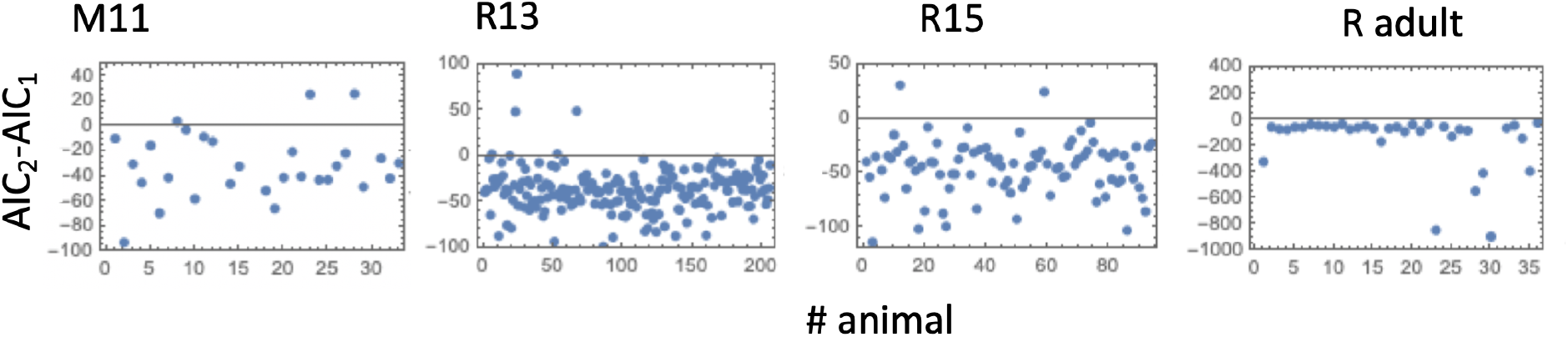
Model selection for the velocity distributions: each dot represents an individual animal and shows the difference *AIC*_2_ − *AIC*_1_, where *AIC*_1_ and *AIC*_2_ are calculated for functions (3) and (4) of the main text respectively. If this quantity is negative, then the model with two terms, function (4), is selected as the more powerful model.

To determine whether the mean values of the two characteristic velocities, *v*_1_ and *v*_2_, were different between different ages, we performed the T-test to compare the group mean of *v*_1_ for R13 with the group mean of *v*_1_ for *R*15 and the group mean of *v*_1_ for R15 with the group mean of *v*_1_ for adult rats, to find that both comparisons yielded a significant difference with *p <* 5 *×* 10^−4^. Similarly, we compared the mean values of *v*_2_ among different groups, and again the T test showed significant differences for all comparisons with *p <* 2 *×* 10^−4^.

## Appendix B The analogy with an alternating-mass Brownian particle

A physical analogy with the obtained velocity dynamics can be constructed in the form of a Brownian particle, whose mass and size randomly vary in time. The mass randomly switches between two values *M*_1_ *> M*_2_, where *M*_1_ corresponds to the slow motion, and *M*_2_ describes the fast movement. This system is equivalent to Brownian motion with fluctuating diffusivity (see [75] and the references therein). The short correlation time *τ*_*s*_ is the time of a single collision with one of the surrounding particles. The large correlation time *τ*_*l*_ is the average time the particle retains a given mass and size before switching to the other mass and size. The friction magnitude and the resulting deceleration are determined by the mass and size.

The probability distribution of the masses is as follows:

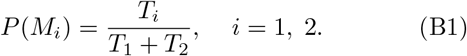

The kinetic energy distribution *E* does not depend on the mass distribution,

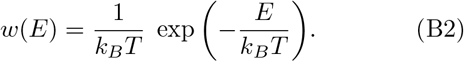

As a result, to obtain their joint distribution, we multiply them, then transform this distribution to the PDF for *v* and *M*_*i*_ and get

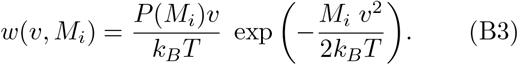

After averaging over *M*_*i*_, we obtain a mixture of two Rayleigh distributions, equation (4) of the main text, where 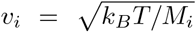 and *P* (*M*_*i*_) = *μ*_*i*_*/*(*μ*_1_ + *μ*_2_). The masses ratio of the two-mode particle should be *M*_1_*/M*_2_ = (*v*_2_*/v*_1_)^2^, and the ratio of the damping coefficients *γ*_1_*/γ*_2_ = (*v*_2_*/v*_1_)^4^. The PDF of the velocity and auto-correlation function are the same as those observed for a rodent.

